# irAEGIS: Pathway-guided interpretable prediction of immune-related adverse events

**DOI:** 10.64898/2026.11.01.763453

**Authors:** Aishwarya Budhkar, Sen Guo, Xuhong Zhang, Jing Su, Qianqian Song

**Affiliations:** Department of Computer Science, Indiana University Bloomington, Indiana, USA; Department of Biostatistics and Health Data Science, Indiana University School of Medicine, Indiana, USA; School of Applied and Creative Computing, Purdue University, Indiana, USA

**Keywords:** Immune-related adverse events (irAE), Single-cell RNA sequencing, Pathway-guided learning, Interpretable deep learning, Phenotype prediction, Biological knowledge integration, Explainable artificial intelligence (XAI)

## Abstract

Immune-related adverse events (irAEs) are major complications of immune checkpoint inhibitor (ICI) therapy and remain difficult to predict prior to treatment initiation. Single-cell RNA sequencing (scRNA-seq) provides a unique opportunity to characterize immune states associated with future irAE development. However, translating single-cell measurements into accurate and biologically interpretable patient-level risk predictions remains challenging. Existing approaches typically aggregate cellular profiles into patient-level summaries, thereby obscuring cell type specific signals, or rely on complex deep-learning architectures that offer limited insight into the biological pathways and mechanisms underlying their predictions. Here, we present irAEGIS, a pathway-guided interpretable model for predicting irAE risk from pre-treatment scRNA-seq data. irAEGIS incorporates prior biological knowledge through gene set constrained representation learning, aggregates pathway activities within individual cell types, and integrates cell type specific risk signals through a hierarchical prediction framework. This design enables predictions to be decomposed into contributions from individual cell types, pathways, and genes, providing interpretable explanations across multiple biological resolutions. Across simulated datasets and independent pre-treatment ICI cohorts, irAEGIS consistently outperformed seven existing methods. Beyond predictive performance, irAEGIS identified biologically coherent immune programs associated with irAE risk. In non-small cell lung cancer, irAEGIS highlighted neutrophil-associated inflammatory pathways linked to future irAE development, whereas in melanoma, it identified apoptotic and immune activation pathways in CD8+ NKT-like cells associated with severe irAEs. These findings were independently supported by pathway enrichment, cell–cell communication, and trajectory analyses. Together, irAEGIS provides a biologically informed and interpretable model for linking single-cell immune states to patient outcomes, enabling both accurate risk prediction and mechanistic insight into immune-related adverse events. More broadly, irAEGIS establishes a general strategy for interpretable phenotype prediction and biological discovery from single-cell transcriptomic data.

**Key Points:**

- irAEGIS is a pathway-guided interpretable framework that predicts immune-related adverse events from pre-treatment single-cell RNA sequencing while providing mechanistic explanations at the cell type, pathway, and gene levels.
- Across simulated datasets and four independent immune checkpoint inhibitor cohorts, irAEGIS consistently outperformed existing machine learning and deep learning approaches for patient-level irAE prediction.
- irAEGIS identified biologically meaningful immune programs associated with irAE risk, including neutrophil-associated inflammatory pathways in non-small cell lung cancer and apoptotic CD8□ NKT-like cell programs associated with severe irAEs in melanoma, which were independently validated by pathway enrichment, cell–cell communication, and trajectory analyses.
- By integrating prior biological knowledge directly into representation learning, irAEGIS establishes a general framework for interpretable phenotype prediction and biological mechanism discovery from single-cell transcriptomic data.

## INTRODUCTION

Single-cell RNA sequencing (scRNA-seq) has transformed the study of complex biological systems by enabling transcriptomic profiling at cellular resolution^1^. Unlike bulk transcriptomic measurements, which average signals across heterogeneous cell populations, scRNA-seq captures diverse cellular states, gene expression programs, and functional heterogeneity within individual tissues and patients^1, 2^. These capabilities have provided unprecedented opportunities to investigate disease mechanisms, therapeutic responses, and clinical outcomes at the resolution of individual cells^3^. As a result, an increasing number of studies now seek to leverage scRNA-seq data not only for descriptive biological analysis but also for predictive modeling of patient phenotypes, including disease progression, treatment response, and adverse clinical outcomes^2, 4^. However, this task presents two fundamental computational challenges. First, predicting patient phenotype from millions of sparse and noisy cell-level measurements requires sophisticated models that can aggregate heterogeneous cellular information into robust patient-level representations^2^. Second, because clinically relevant phenotypes often arise from specific cell populations and biological programs, effective prediction models should also provide interpretable explanations that identify the cell types, pathways, and genes driving each prediction. Such interpretability can increase trust in model predictions and help uncover biologically meaningful mechanism.

Immune checkpoint inhibitors (ICIs) have revolutionized cancer treatment by restoring anti-tumor immune activity and producing durable clinical responses across multiple malignancies^5^. Despite their clinical success, ICIs can trigger immune-related adverse events (irAEs), a broad spectrum of inflammatory toxicities that can affect nearly every organ system. irAEs range from mild symptoms requiring minimal intervention to severe and potentially life threatening complications that may necessitate treatment discontinuation or long-term immunosuppression^5, 6^. Consequently, identifying patients at elevated risk of developing irAEs before treatment initiation remains an important unmet clinical need. Because irAEs arise from dysregulated immune activation, pre-treatment profiling of peripheral immune cells provides a promising strategy for identifying molecular and cellular programs associated with future toxicity^6^. In particular, scRNA-seq of peripheral blood mononuclear cells (PBMCs) enables comprehensive characterization of immune-cell composition, activation states, and functional programs that may predispose patients to irAE development^5^.

Recent advances in machine learning and deep learning have enabled increasingly sophisticated approaches for phenotype prediction from scRNA-seq data^7, 8^. Some machine learning approaches often aggregate cellular profiles into patient-level pseudobulk representations. For example, pseudobulk logistic regression^9^ uses aggregated patient-level gene expression features with a linear classifier (e.g. Random forest^10^, XGBoost^11^) for binary outcome prediction. While computationally efficient, they lose information regarding cellular heterogeneity and may obscure signals originating from rare but biologically important cellular subpopulations^12^. Some deep learning methods have also been proposed for phenotype prediction from scRNA-seq data. For example, ScRAT^13^ uses a transformer-based attention mechanism with mixup augmentation to identify informative cells for patient phenotype prediction. SingleDeep^14^ uses adaptive neural network architectures for sample phenotype prediction and utilizes the post-hoc feature attribution method DeepLIFT^15^ to estimate gene contributions within different cell types. Although, this provides explanations, post-hoc attributions can be sensitive to the attribution method and the input gene expression, making them difficult to align directly with established biological pathways^16, 17^. Hierarchical MIL^18^ leverages multiple-instance learning with hierarchical attention to aggregate information across cells and cell types for patient phenotype prediction. However, its interpretability is primarily based on attention-derived importance rather than biological knowledge. Thus, while these methods improve the modeling of cellular heterogeneity for effective translation of single-cell measurements into patient-level predictions, their predictions are often generated through complex latent representations and attention mechanisms that are difficult to map back to biologically meaningful pathways, cellular programs, or molecular mechanisms^16, 19^.

This limitation is particularly important in clinical settings, where biological interpretability is often as important as predictive accuracy. For irAE prediction, a useful model should not only estimate whether a patient is likely to develop toxicity but also indicate which immune cell populations and molecular pathways contribute to that risk. Such interpretation is essential for generating biological hypotheses, identifying potential biomarkers, and understanding disease mechanisms^20^. We therefore hypothesized that incorporating prior biological knowledge directly into representation learning could help improve both patient-level phenotype prediction and biological interpretability. In this work, we propose irAEGIS, a pathway-guided interpretable framework for predicting irAE risk from scRNA-seq data before immunotherapy. irAEGIS incorporates prior biological knowledge through gene set constrained representation learning, aggregates pathway activities within individual cell types, and integrates cell type specific risk signals through a hierarchical prediction framework. By performing prediction in biologically meaningful pathway space rather than directly in high-dimensional gene space, irAEGIS enables predictions to be decomposed into contributions from individual cell types, pathways, and genes. We evaluated irAEGIS across simulated datasets and multiple independent pre-treatment ICI cohorts and compared its performance against seven existing methods. Beyond achieving superior predictive performance, irAEGIS identified biologically coherent immune programs associated with irAE risk, including neutrophil-associated inflammatory pathways and CD8+ NKT-like apoptotic programs that were independently supported by pathway enrichment, cell-cell communication, and trajectory analyses. More broadly, irAEGIS establishes a general framework for interpretable phenotype prediction and biological discovery from single-cell transcriptomic data.

## RESULTS

### Overview of the irAEGIS model

The primary goal of irAEGIS is to predict immune-related adverse event (irAE) risk from pre-treatment single-cell transcriptomic data while providing biologically interpretable explanations across multiple levels of cellular organization. **Figure 1** illustrates the overall workflow of irAEGIS. The framework takes pre-treatment PBMC scRNA-seq profiles together with cell type annotations as input and generates both patient-level irAE risk predictions and interpretable biological insights.

**Figure 1:**
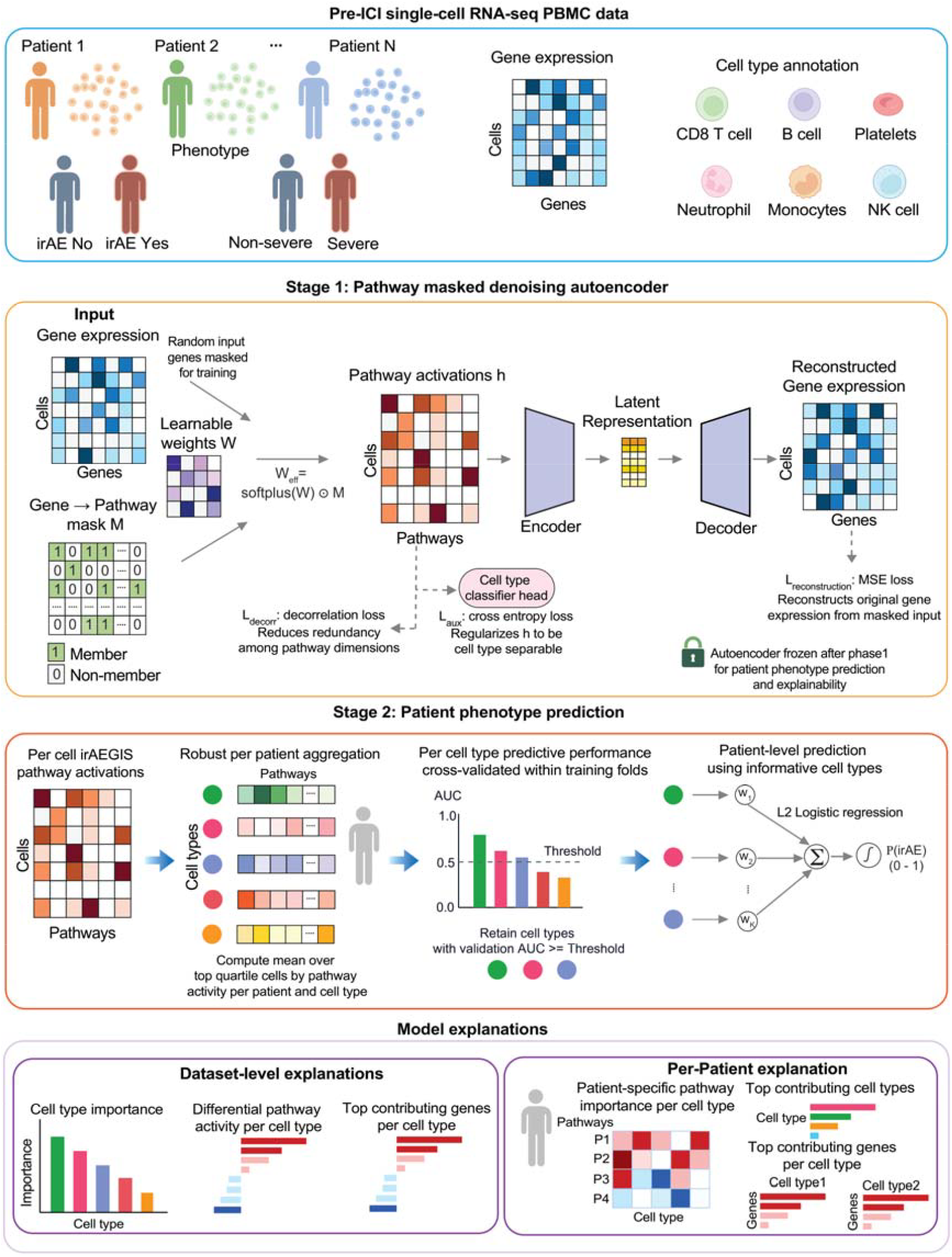
Overview of the irAEGIS model. **(a)** irAEGIS uses pre-treatment PBMC scRNA-seq data, cell type annotations, and patient irAE label to predict irAE risk and generate biologically resolved explanations. **(b)** Pathway masked denoising autoencoder: irAEGIS randomly masks input gene expression and uses a Hallmark gene to pathway mask to produce pathway activity representations in which each dimension corresponds to a predefined biological program. The autoencoder maps the masked input into low-dimensional latent representation and reconstructs the original unmasked gene expression profile, while jointly preserving cell type identity and encouraging pathway activations to be decorrelated. Next, the autoencoder is frozen for downstream prediction. **(c)** Patient phenotype prediction: irAEGIS aggregates pathway representations within each patient and cell type by averaging cells in the upper quartile of overall pathway activity. Cell type specific classifiers are then trained to estimate irAE risk, and informative cell types selected by inner cross validation AUC are integrated to produce final patient-level prediction. **(d)** irAEGIS provides explanations at both dataset and patient-level. Dataset-level explanations include cell type importance ranking, differential pathway activity, and top contributing genes per cell type. Patient-level explanations identify the cell types, pathways, and genes per cell type contributing to an individual prediction.

irAEGIS consists of two major stages. In the first stage, individual cells are encoded into biologically meaningful pathway representations through a pathway-guided denoising autoencoder. Rather than learning latent features directly from gene expression alone, the encoder incorporates prior biological knowledge by constraining representations to predefined gene sets, allowing each dimension to capture the activity of a specific biological program. The autoencoder is trained to reconstruct the original gene expression profile while preserving cell type information, thereby learning pathway representations that are both biologically grounded and robust to technical noise. In the second stage, cell-level pathway activities are aggregated within each patient and cell type to generate cell type resolved pathway profiles. This aggregation strategy emphasizes highly activated cellular subpopulations that may carry phenotype-associated signals while preserving cellular heterogeneity across immune populations. Cell type specific classifiers are then trained to estimate irAE risk independently within each immune cell population. Finally, a patient-level classifier integrates the risk signals from informative cell types to generate a unified patient-level prediction. At inference, the trained encoder maps cells from previously unseen patients into pathway space, after which pathway activities are aggregated and passed through the cell type specific and patient-level classifiers to produce an irAE risk score. Importantly, because prediction is performed using biologically interpretable pathway representations, the framework naturally supports mechanistic interpretation. Importantly, irAEGIS provides explanations at both the dataset and patient levels. At the dataset level, the framework identifies the immune cell types, biological pathways, and genes most strongly associated with irAE risk across a cohort. At the individual patient-level, irAEGIS quantifies how specific cell types, pathways, and genes contribute to a prediction, enabling mechanistic interpretation of patient-specific risk. Together, these capabilities allow irAEGIS to move beyond black-box prediction by linking patient outcomes to the underlying cellular and molecular programs that drive them.

### irAEGIS achieves robust predictive performance and recovers known biological signals in simulated datasets

To evaluate both predictive performance and biological interpretability under controlled conditions, we benchmarked irAEGIS against seven representative methods using two complementary simulation settings with known ground-truth signals. The distributed signal (DS) setting models a scenario in which weak phenotype-associated signals are distributed across multiple immune cell types, whereas the restricted signal (RS) setting represents a more localized scenario in which stronger signals are confined to a subset of cell populations. We compared irAEGIS against three deep-learning methods for single-cell phenotype prediction (ScRAT, SingleDeep, and Hierarchical MIL) and four traditional machine-learning baselines (Cell LR, pseudobulk Logistic Regression (LR), Random Forest, and XGBoost). Performance was evaluated using the area under the receiver operating characteristic curve (AUC) and area under the precision–recall curve (AUPRC) computed from out-of-fold patient predictions.

Across both simulation settings, irAEGIS consistently achieved the best predictive performance **(Figure 2a-b; Supplementary Table S1)**. In the DS setting, irAEGIS achieved the highest average AUC (0.849) and AUPRC (0.862), substantially outperforming all competing methods. Similar results were observed in the RS setting, where irAEGIS achieved an average AUC of 0.854 and AUPRC of 0.831, again exceeding all baseline approaches. Notably, irAEGIS maintained strong performance in both distributed and cell type restricted signal settings, indicating robustness to different biological architectures of disease-associated signals. We next evaluated whether the features prioritized by irAEGIS were biologically meaningful **(Figure 2c)**. Fidelity analysis showed that masking genes belonging to the top-ranked irAEGIS pathways substantially reduced predictive performance, with median AUC decreases of 0.223 and 0.411 in the DS and RS settings, respectively. Similarly, masking the informative cell types led to marked reductions in prediction accuracy. These results indicate that the pathways and cell types identified by irAEGIS are functionally important for model predictions rather than incidental correlates.

**Figure 2:**
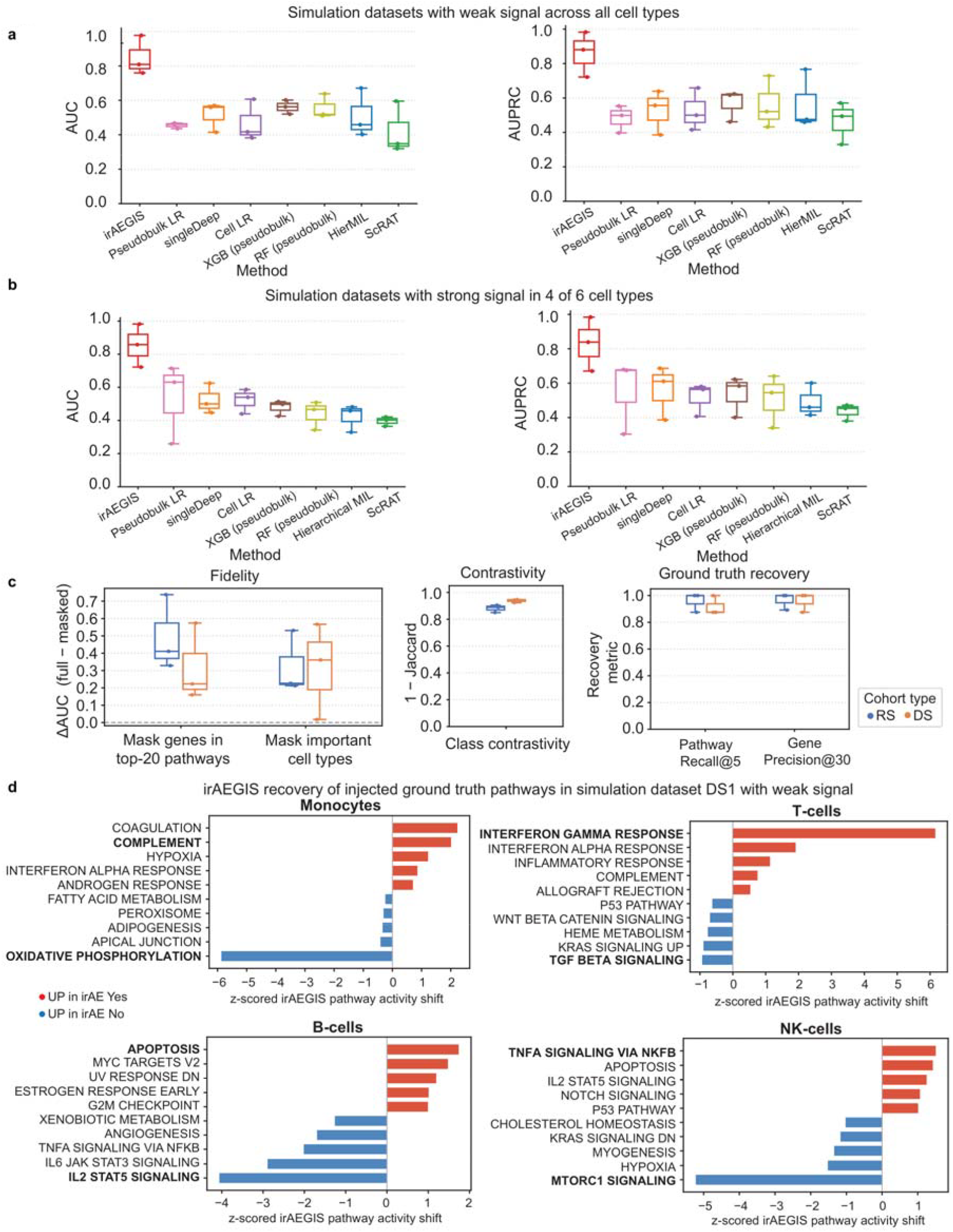
Evaluation of predictive performance and biological signal recovery in simulated datasets, **(a)** Box plots of AUC and AUPRC for each method across three DS simulated datasets, in which weak irAE signal is distributed across all four simulated immune cell types. **(b)** Box plots of AUC and AUPRC for each method across three RS simulated datasets, in which stronger irAE signal is restricted to four of six simulated cell types. **(c)** Box plots summarizing pathway and cell type fidelity, class contrastivity, and ground truth gene and pathway recovery for irAEGIS across simulated datasets. **(d)** Recovery of injected ground truth pathways across the four simulated immune cell types in the DS1 dataset. Bars show z-scored differential pathway activity between irAE positive and negative groups for the irAEGIS important pathways within each cell type. Red indicates higher activity in irAE positive patients, and blue indicates higher activity in irAE negative patients. Injected ground truth pathways are highlighted in bold.

To assess whether irAEGIS distinguishes biologically meaningful programs between phenotype groups, we evaluated class contrastivity. irAEGIS achieved median contrastivity scores of 0.944 in the DS setting and 0.889 in the RS setting, indicating that the pathways associated with irAE positive and irAE negative patients were largely distinct. We further quantified recovery of the known ground-truth signals embedded in the simulations. irAEGIS recovered 87.5-100% of the injected pathways among the top-ranked pathways for each signal-carrying cell type and achieved 87.5-100% overlap between the top-ranked genes and the injected ground-truth gene sets. These results demonstrate that the biological explanations generated by irAEGIS accurately reflect the underlying mechanisms used to construct the simulated datasets. **Figure 2d** illustrates pathway recovery in the DS dataset. Across all four simulated immune cell types, irAEGIS successfully recovered the injected pathways together with their expected directions of regulation. For example, irAEGIS identified increased COMPLEMENT activity in irAE positive monocytes and increased Oxidative Phosphorylation activity in irAE negative monocytes. Similarly, it correctly recovered Interferon Gamma Response and TGF-β Signaling programs in T cells, Apoptosis and IL2-STAT5 Signaling programs in B cells, and TNF-*α* Signaling via NF-B and mTORC1 Signaling programs in NK cells. The recovered pathway patterns closely matched the known simulation design, demonstrating that irAEGIS can accurately localize phenotype-associated signals to the correct cell types and biological programs. Together, these results demonstrate that irAEGIS not only achieves superior predictive performance but also faithfully recovers biologically meaningful signals at the cell type, pathway, and gene levels. This ability to jointly optimize prediction and biological interpretability provides the foundation for subsequent analyses in real patient cohorts.

### irAEGIS identifies biologically coherent immune programs associated with irAE risk in real patient cohorts

Having established that irAEGIS accurately recovers known biological signals in simulated datasets, we next evaluated its predictive performance and interpretability in real-world patient cohorts. We benchmarked irAEGIS against seven representative methods across four independent pre-treatment immune checkpoint inhibitor (ICI) datasets: GSE189125, GSE216329, GSE249898, and GSE285888 (**Supplementary Table S1**). Model performance was assessed using patient-level AUC and AUPRC computed from held-out predictions across cross-validation folds.

Across all four cohorts, irAEGIS consistently achieved the strongest predictive performance **(Figure 3a)**. The model obtained the highest average AUC (0.79) and AUPRC (0.763), outperforming all competing approaches, including both pseudobulk machine learning methods and recent deep learning models for single-cell phenotype prediction. These results demonstrate that biologically informed pathway representations can improve prediction across diverse clinical cohorts while preserving cellular resolution. We next examined whether the biological explanations generated by irAEGIS remained informative in real patient datasets. Fidelity analysis showed that masking genes belonging to the top-ranked irAEGIS pathways resulted in a median AUC decrease of 0.456, whereas masking the most informative cell types produced a median AUC decrease of 0.494 **(Figure 3b)**. These results indicate that the pathways and cell types prioritized by irAEGIS contribute substantially to model predictions. Furthermore, irAEGIS achieved a median class-contrastivity score of 0.748, suggesting that the pathway programs associated with distinct irAE phenotypes are largely separable and biologically distinct across cohorts.

**Figure 3:**
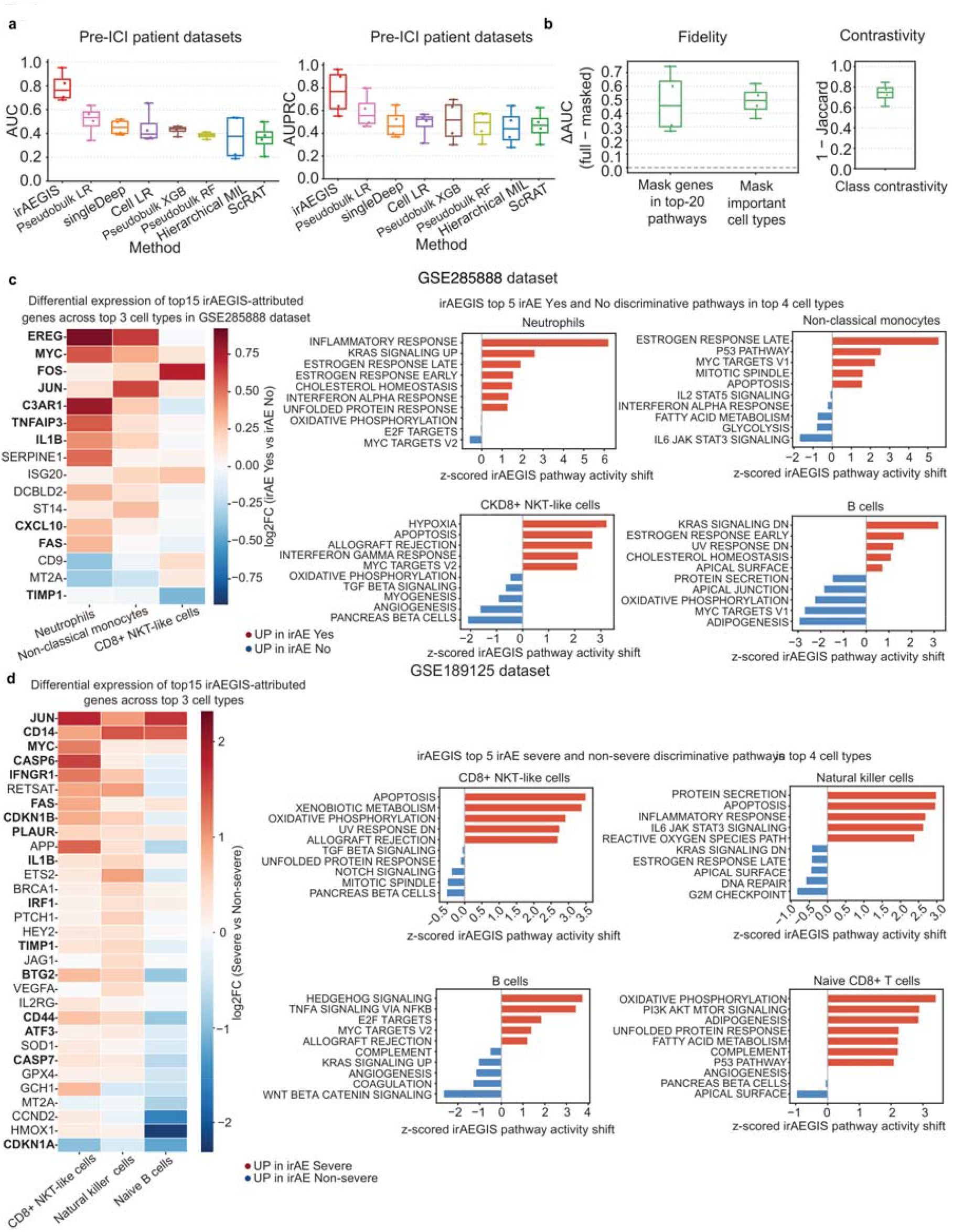
Evaluation of predictive performance and model derived biological explanations in real patient datasets. **(a)** Box plots of AUC and AUPRC across four pre-treatment immune checkpoint inhibitor (ICI) datasets: GSE189125, GSE216329, GSE249898, and GSE285888, comparing irAEGIS with seven competing methods. **(b)** Box plots of pathway and cell type fidelity, and class contrastivity for irAEGIS across the four real datasets. **(c)** irAEGIS explanations for the GSE285888 NSCLC dataset. Left: heatmap of log2 fold-change between positive and negative irAE patients for the top 15 irAEGIS important genes across the three informative cell types. Right: top five Hallmark pathways associated with irAE positive and irAE negative status across the top ranked cell types. **(d)** irAEGIS explanations for the GSE189125 melanoma dataset. Left: heatmap of log2 fold-change between severe and non-severe irAE patients for the top 15 irAEGIS important genes across the three informative cell types. Right: top five Hallmark pathways associated with severe and non-severe irAE status across the top ranked cell types.

We then investigated the biological programs prioritized by irAEGIS in individual datasets. In the GSE285888 NSCLC cohort, irAEGIS identified neutrophils, non-classical monocytes, and CD8+ NKT-like cells as the most informative cell populations associated with future irAE development **(Figure 3c)**. Within these cell types, genes including *EREG, MYC, FOS, JUN, C3AR1, TNFAIP3, IL1B, ISG20, CXCL10*, and *FAS* were more highly expressed in irAE positive patients, whereas *CD9, MT2A*, and *TIMP1* were associated with irAE negative patients. At the pathway level, Inflammatory Response, Apoptosis, and Interferon Response programs were among the strongest contributors to irAE risk. These findings are consistent with previous studies implicating inflammatory myeloid states and neutrophil-associated immune activation in ICI-induced toxicity^21-23^. We next analyzed the GSE189125 melanoma cohort, in which patients were stratified according to severe versus non-severe irAE phenotypes **(Figure 3d)**. irAEGIS identified CD8+ NKT-like cells, natural killer cells, and naïve B cells as the most informative cell populations. Genes including *JUN, CD14, MYC, CASP6, IFNGR1, FAS, CDKN1B, PLAUR, IL1B, IRF1, BTG2, CD44, ATF3*, and *CASP7* were preferentially associated with severe irAE, whereas *CDKN1A, MT2A, GCH1*, and *CCND2* were enriched in non-severe patients. Consistent with these gene-level patterns, Apoptosis, Allograft Rejection, and Inflammatory Response pathways were strongly associated with severe irAE. These observations agree with previous reports linking pre-treatment cytotoxic T-cell and NK-cell programs to severe immune-related toxicity. Together, these results demonstrate that irAEGIS not only improves predictive performance across independent patient cohorts but also identifies biologically coherent cell type specific programs associated with irAE development and severity. The consistency between model-derived explanations and previously reported irAE mechanisms supports the biological validity of the pathway, cell type, and gene-level insights generated by irAEGIS.

### irAEGIS identifies a neutrophil-associated inflammatory program linked to irAE development

To evaluate the biological relevance of the signals identified by irAEGIS, we performed a detailed analysis of the GSE285888 NSCLC cohort. We used the cell type, pathway, and gene-level explanations generated by irAEGIS to investigate immune programs associated with future irAE development and then assessed these findings using independent downstream analyses.

Among the cell populations prioritized by irAEGIS, neutrophils exhibited strong transcriptional differences between irAE positive and irAE negative patients as reflected by the top ranked neutrophil genes in **Figure 4a**. Several of these genes, including *CCL20, EREG, C3AR1, CSF1, MYC, IL1B, CXCL10*, and *IRF1* showed higher expression in irAE positive patients and are involved in inflammatory, cytokine, and chemokine signaling. These observations suggest the presence of a pre-treatment inflammatory neutrophil program associated with future irAE development and are consistent with previous studies linking activated neutrophils and inflammatory myeloid states to ICI-associated toxicity^21-23^. To independently evaluate the pathway-level interpretations produced by irAEGIS, we performed gene set enrichment analysis (GSEA) using differential expression between irAE positive and irAE negative patients. Consistent with the model-derived explanations, the Hallmark Inflammatory Response pathway was significantly enriched in irAE positive patients (NES = 1.59, FDR = 0.0038; **Figure 4b**). This pathway was also identified by irAEGIS as one of the most important programs within neutrophils. More broadly, 70% of pathways showed concordant directions of regulation between irAEGIS-derived pathway activity shifts and enrichment scores through differential expression analysis, supporting the robustness and biological validity of the pathway representations learned by the model.

**Figure 4:**
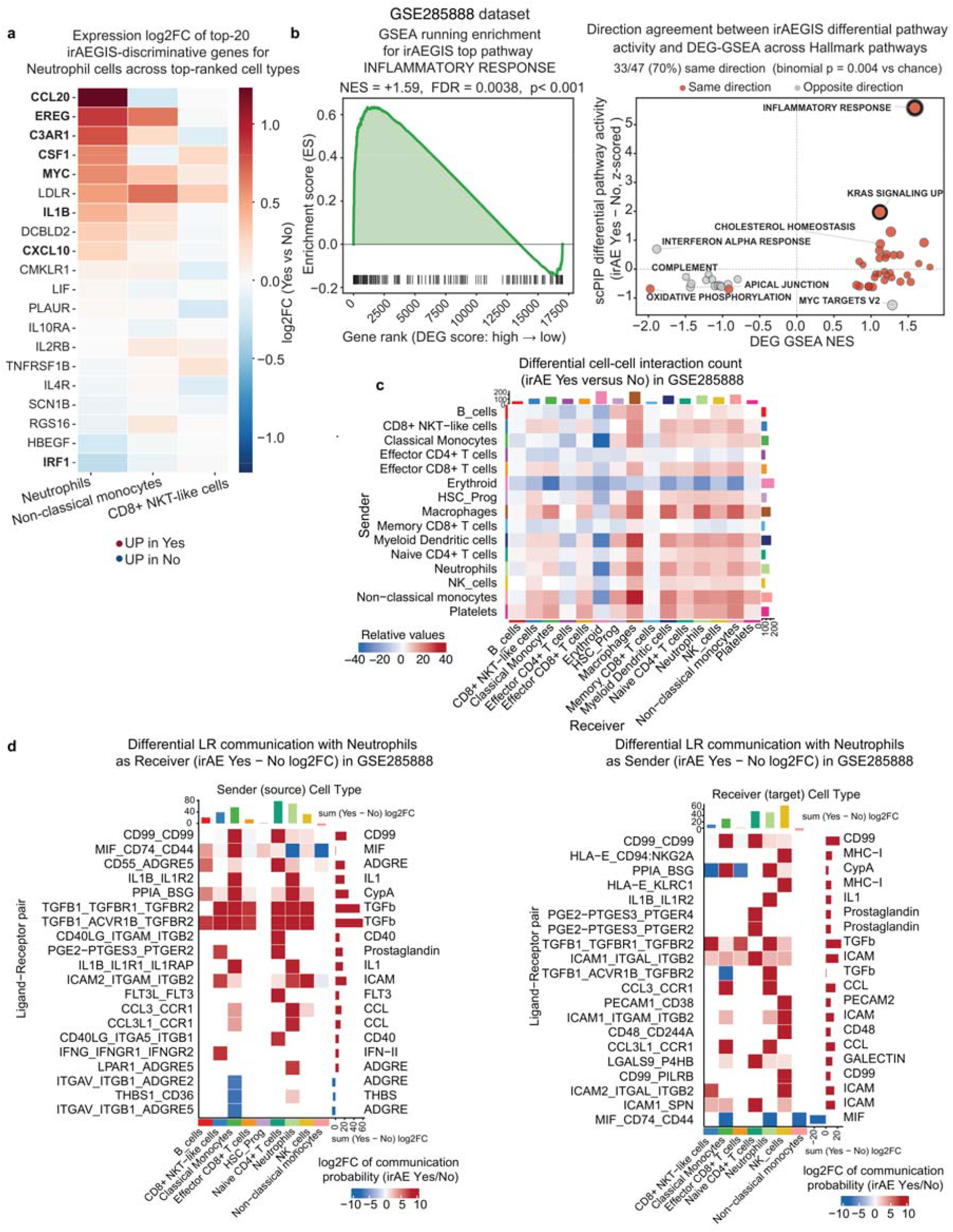
Neutrophil associated inflammatory program identified by irAEGIS for distinguishing irAE positive and irAE negative patients. **(a)** Heatmap of log2 fold-change between irAE positive and negative patients for the top 20 irAEGIS important genes in neutrophils. **(b)** Independent evaluation of irAEGIS pathway explanations using GSEA. Left: GSEA running enrichment score for the Hallmark Inflammatory Response gene set based on differential expression between irAE positive and irAE negative patients. **(c)** Differential cell-cell interaction counts between irAE positive and irAE negative patients across all cell type pairs in GSE285888 inferred using CellChat analysis. **(d)** Differential ligand-receptor communication involving neutrophils as receivers on the left and as senders on the right in irAE positive versus irAE negative patients.

We then examined whether the inflammatory neutrophil program identified by irAEGIS was accompanied by altered intercellular communication. To address this question, we applied CellChat^24^ to infer ligand-receptor interactions and compared communication networks between irAE positive and irAE negative patients. Neutrophils exhibited increased interactions with monocytes, macrophages, and myeloid dendritic cells in irAE positive patients **(Figure 4c)**, suggesting the presence of a coordinated inflammatory communication network prior to treatment initiation. These observations are consistent with reports linking baseline inflammatory immune states and elevated cytokine signaling to subsequent irAE development^25, 26^. To further characterize these communication changes, we examined the ligand-receptor interactions involving neutrophils as both signal receivers and senders **(Figure 4d)**. As receivers, neutrophils displayed increased incoming signaling through IL1, TGFβ, CD40, prostaglandin (*PGE2*), and CCL pathways, primarily originating from classical monocytes, CD8+ NKT-like cells, naive CD4+ T cells, and other neutrophils. As senders, neutrophils showed increased outgoing signaling toward natural killer cells, naive CD4+ T cells, and neighboring neutrophils through MHC-I, IL1, ICAM, prostaglandin, and CCL-associated interactions. Notably, several prioritized interactions have previously been implicated in immune regulation and ICI-associated toxicity, including HLA-E–CD94:NKG2A signaling and IL-1β mediated inflammatory pathways^23, 27, 28^. Together, these analyses provide convergent evidence from differential expression, pathway enrichment, and cell-cell communication analyses that the signals prioritized by irAEGIS reflect a biologically coherent neutrophil-centered inflammatory program associated with future irAE development. These findings support the interpretability of the model and demonstrate its ability to uncover mechanistically meaningful immune programs from pre-treatment single-cell transcriptomic data.

### irAEGIS reveals a CD8+ NKT-like apoptotic program associated with severe irAE

We next investigated whether the biological signals prioritized by irAEGIS could reveal immune programs associated with severe irAE phenotypes, using the GSE189125 melanoma cohort. Leveraging the cell type, pathway, and gene-level explanations generated by irAEGIS, we examined transcriptional programs that distinguish patients who developed severe irAEs from those with non-severe toxicity and evaluated these findings using independent downstream analyses.

At the gene level, irAEGIS highlighted a transcriptional program enriched for apoptosis, inflammatory signaling, cytokine response, and immune regulation **(Figure 5a)**. Among the top-ranked genes associated with severe irAE were *JUN, CASP6, FAS, CDKN1B, GCH1, PLAUR, CASP7, TIMP1*, and *HMOX1*, many of which are involved in cell death, stress responses, and immune activation. These observations are consistent with previous studies reporting associations between baseline cytotoxic T-cell and NK-cell states and subsequent irAE development^5, 29-31^. To independently evaluate the pathway-level explanations produced by irAEGIS, we performed gene set enrichment analysis (GSEA) using differential expression between severe and non-severe irAE patients. Consistent with the model-derived results, the Hallmark Apoptosis pathway was significantly enriched in severe irAE patients (NES = 1.26, FDR = 0.0034; **Figure 5b**). Across all Hallmark pathways, 87% showed concordant directions between irAEGIS-derived pathway activity shifts and enrichment scores through differential expression analysis, supporting the reproducibility and biological validity of the pathway representations learned by the model.

**Figure 5:**
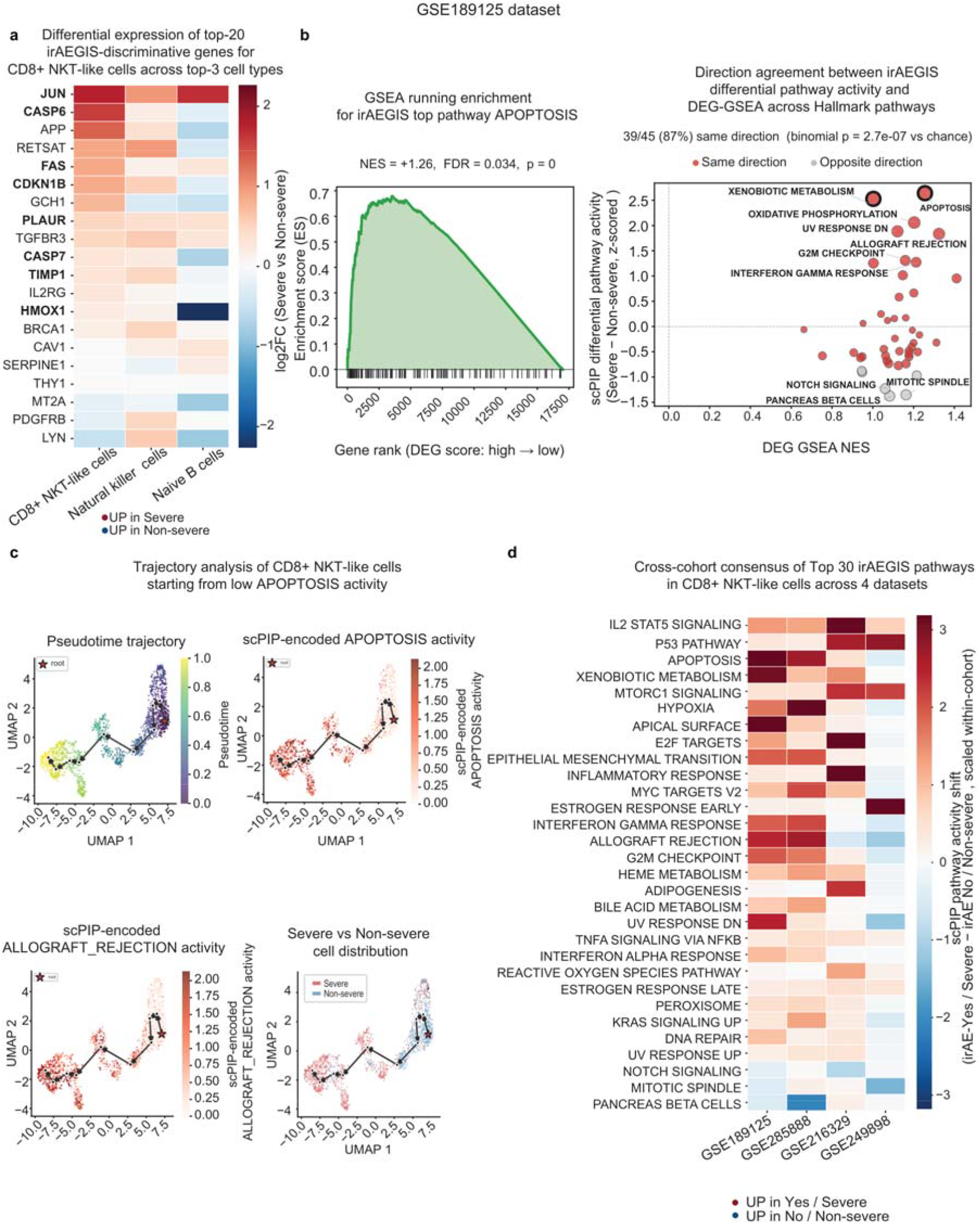
CD8+ NKT-like apoptotic program identified by irAEGIS for distinguishing severe and non-severe irAE patients. **(a)** Heatmap of log2 fold-change between severe and non-severe irAE patients for the top 20 irAEGIS important genes in CD8+ NKT-like cells. **(b)** Independent evaluation of irAEGIS pathway explanations using GSEA. Left: GSEA running enrichment score for the Hallmark Apoptosis gene set based on differential expression between severe and non-severe irAE patients. **(c)** Pseudotime trajectory analysis of CD8+ NKT-like cells. Top left: UMAP projection colored by inferred pseudotime, with the trajectory rooted at cells with low apoptosis gene set activity from raw gene expression. Top right: UMAP projection colored by irAEGIS encoded Apoptosis pathway activity. Bottom left: UMAP projections colored by irAEGIS encoded Allograft Rejection pathway activity. Bottom right: UMAP projection colored by severity group, with cells from severe irAE patients shown in red and cells from non-severe irAE patients shown in blue. **(d)** Cross dataset consensus heatmap of the top 30 irAEGIS important Hallmark pathways for CD8+ T /NK populations across datasets. Each heatmap entry shows normalized irAEGIS differential pathway activity, defined as the mean pathway activity difference between irAE groups, scaled by the standard deviation of pathway shifts in that dataset. This preserves the direction of change while enabling comparison across datasets. Red indicates higher activity in irAE positive or severe patients, and blue indicates higher activity in irAE negative or non-severe patients.

We then evaluated whether the pathways prioritized by irAEGIS were associated with cellular state transitions. To address this question, we inferred pseudotime trajectories within CD8+ NKT-like cells using Monocle3^32^ and projected irAEGIS-derived pathway activities onto the resulting trajectory **(Figure 5c)**. Apoptosis activity increased progressively along pseudotime, and a similar trend was observed for the Allograft Rejection pathway. Mapping patient severity labels onto the same trajectory revealed that cells from severe irAE patients were preferentially enriched toward the high-activity end of the trajectory, whereas cells from non-severe patients were concentrated in regions with lower pathway activity. These results suggest that the biological programs prioritized by irAEGIS correspond to continuous cellular state transitions associated with severe toxicity risk. To determine whether these findings were specific to GSE189125 or reflected broader immune programs associated with irAE, we compared the top pathways prioritized by irAEGIS across all four pre-treatment ICI cohorts. Several pathways, including IL2-STAT5 Signaling, mTORC1 Signaling, P53 Pathway, Xenobiotic Metabolism, Apoptosis, and MYC Targets V2, were consistently elevated in irAE positive or severe-irAE groups across multiple datasets **(Figure 5d)**. The recurrence of these pathways across independent cohorts suggests that the biological programs identified by irAEGIS represent reproducible features of pre-treatment immune states associated with irAE susceptibility. Together, these analyses provide convergent evidence from differential expression, pathway enrichment, pseudotime analysis, and cross-cohort comparison that the signals prioritized by irAEGIS reflect biologically meaningful immune programs associated with severe irAE. These findings further support the interpretability of the model and demonstrate its ability to uncover reproducible cellular mechanisms linked to immune-related toxicity.

## DISCUSSION

Single-cell transcriptomics has enabled unprecedented characterization of cellular heterogeneity across health and disease. However, most biological and clinical questions are ultimately defined at the patient-level rather than the cellular level. Treatment response, disease progression, toxicity risk, and clinical outcomes are all patient-level phenotypes that emerge from the collective behavior of diverse cellular populations. A fundamental challenge therefore lies in connecting cellular states measured by single-cell technologies to clinically relevant patient phenotypes while preserving biological interpretability. In this study, we developed irAEGIS to address this challenge. Rather than viewing phenotype prediction solely as a machine learning task, irAEGIS provides a biologically informed framework that links cell type specific molecular programs to patient-level outcomes. Rather than treating prediction and interpretation as separate tasks, irAEGIS integrates prior biological knowledge into representation learning so that predictive signals are organized around biologically meaningful pathway activities within individual cell types. This design enables patient-level predictions to be decomposed into contributions from specific cell populations, molecular pathways, and genes, providing a direct connection between cellular programs and clinical phenotypes. Across simulated datasets and independent pre-treatment

ICI cohorts, irAEGIS consistently outperformed existing methods while maintaining biological interpretability. Furthermore, simulation studies demonstrated that the framework accurately recovered known cell type, pathway, and gene-level signals, indicating that the explanations generated by the model faithfully reflect the underlying biology.

Beyond predictive performance, our results suggest that interpretable representation learning can help connect patient-level risk prediction to biologically meaningful immune programs. In the GSE285888 cohort, irAEGIS identified a neutrophil-associated inflammatory program, which is consistent with prior reports linking activated neutrophils, inflammatory response pathways, enhanced communication with other myeloid populations, and baseline cytokine signaling to ICI associated toxicity and subsequent irAE development^21, 25, 26, 33, 34^. In the GSE189125 cohort, irAEGIS identified apoptosis and immune activation associated programs in CD8+ NKT-like cells, aligning with studies that implicate pre-treatment cytotoxic T cell states, CD8 T cell populations, NK cell populations, and altering circulating T cell compartments in irAE incidence or severity^29-31, 34^. Importantly, the consistency of these model derived programs with pathway enrichment, cell-cell communication, pseudotime, and cross-cohort analyses supports the biological plausibility of the explanations generated by irAEGIS. Together, these results suggest that irAEGIS can move beyond patient-level risk scoring and biomarker identification to reveal candidate cellular programs and immune mechanisms associated with disease phenotypes.

Of note, this study highlights the importance of incorporating biological knowledge directly into predictive models to bridge cellular states and patient-level phenotypes. Most existing explainable AI approaches attempt to interpret predictions after model training through post-hoc analyses, often making it difficult to determine how learned features relate to underlying biological mechanisms. In contrast, irAEGIS integrates biological priors directly into representation learning, organizing cellular information around biologically meaningful pathway activities from the outset. As a result, the learned representations are not only predictive but also mechanistically interpretable, enabling patient-level phenotypes to be linked to specific cell types, molecular pathways, and genes. We believe this capability is particularly important for single-cell studies. Although single-cell technologies provide unprecedented resolution for characterizing cellular heterogeneity, many biologically and clinically relevant questions, including treatment response, disease progression, toxicity risk, and disease severity, are ultimately defined at the patient level. Consequently, a major challenge is not simply predicting patient outcomes from single cell data but understanding how cellular programs collectively give rise to those outcomes. By connecting cell type specific pathway activities to patient phenotypes, irAEGIS provides a general framework for linking cellular heterogeneity to clinical phenotypes and their underlying biological mechanisms. Although developed for immune-related adverse event prediction, the conceptual framework underlying irAEGIS is broadly applicable. Many clinically relevant phenotypes arise from dysregulation of specific biological programs within distinct cellular populations, including treatment response, therapeutic resistance, autoimmune disease activity, infection severity, and cancer progression. The ability to identify phenotype-associated cell types, pathways, and genes within a unified predictive framework may facilitate both biomarker discovery and mechanistic investigation across a wide range of biomedical applications.

Despite these advantages, several limitations should be considered. First, irAEGIS relies on accurate cell type annotations, and annotation errors may affect downstream interpretations. Second, the current framework focuses exclusively on transcriptomic measurements and does not incorporate complementary modalities such as proteomics, spatial transcriptomics, histopathology, or clinical variables, which may further improve both predictive performance and biological characterization. Third, although irAEGIS can accommodate diverse pathway databases, any predefined gene set collection provides only a partial representation of cellular biology and may fail to capture previously uncharacterized biological programs. Finally, the cellular mechanisms identified in this study should be regarded as computationally generated hypotheses that require further experimental and clinical validation. In summary, irAEGIS establishes a biologically informed framework that directly links single-cell states to patient-level phenotypes through interpretable pathway representations. Beyond improving predictive performance, the framework enables mechanistic interpretation of patient outcomes and provides a generalizable strategy for phenotype prediction, biomarker discovery, and biological mechanism identification from single-cell transcriptomic data.

## MATERIALS AND METHODS

### Data processing

For performance evaluation, we used four pre-treatment ICI datasets that profiled PBMCs by single-cell RNA sequencing: GSE189125 (melanoma), GSE216329 (non-small cell lung cancer (NSCLC), head and neck squamous cell carcinoma (HNSCC), prostate cancer), GSE249898 (not specified), and GSE285888 (NSCLC). We also evaluated irAEGIS on two simulated settings with known ground truth signal. The restricted signal setting (RS) contains irAE discriminative signal injected into four of six simulated immune cell types, and the distributed signal setting (DS) contains weaker irAE discriminative signal distributed across all four simulated cell types. A summary of datasets is provided in **Supplementary Table S2**. For each real cohort, cells expressing fewer than 200 genes were removed to reduce noise from low quality cells. To support stable model training, cell types with less than 50 cells or less than two patients in any irAE status group were excluded. Gene expression was first normalized to 10000 counts per cell and then log transformed to stabilize variance and reduce data skewness. The final gene set included all genes represented in MSigDB^35^ Hallmark gene sets along with the top 2000 highly variable genes. Patient irAE labels were derived from the metadata of each dataset. For GSE189125 and GSE249898, patients with irAE grade 3 or higher were labeled as severe. Cell type annotation was performed using scType^36^. The simulated datasets were generated using Splatter^37^ R package, with three batches of 30 patients each. irAE signals were injected through Hallmark pathways^35^ at varying fold change scales across batches, allowing evaluation against known cell type, pathway, and gene ground truth.

### The irAEGIS model

irAEGIS consists of a pathway-constrained denoising autoencoder with an auxiliary cell type specific classification objective, cell type specific classifiers, and a patient-level classifier that integrates cell type risk scores into a final irAE prediction. The autoencoder follows the denoising autoencoder framework^38^, in which the encoder maps a randomly masked gene expression profile to a low-dimensional representation and the decoder reconstructs the original expression profile. This design encourages the encoder to learn representations that are robust to technical noise and dropout. The encoder is a multi-layer fully connected neural network and includes a pathway activity layer constrained by Hallmark gene set mask, such that each dimension represents a distinct biological program. During training, the autoencoder is also optimized to predict the cell type of each cell from the pathway activity representation, encouraging the learned pathway space to preserve the cell type variation needed for downstream cell type modeling and attribution. After autoencoder training, pathway representations are aggregated within each patient and cell type. Cell type specific classifiers then generate patient-level irAE scores from these aggregated representations. Finally, the patient-level classifier integrates the scores from informative cell type specific classifiers to produce the final patient-level irAE prediction.

### Autoencoder

The irAEGIS autoencoder consists of an encoder *E* and a decoder *D* . The encoder maps an input gene expression vector *x* ∈ ℝ^*G*^ to a low-dimensional latent representation *z* ∈ ℝ^*L*^ through an intermediate pathway activity representation *h* ∈ ℝ^*P*^. The decoder *D* reconstructs gene expression 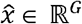 from *z*.

During training, each input expression vector was corrupted by random gene masking as follows:

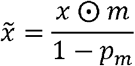

Where *m* ∈ {0,1}^*G*^ is a binary mask sampled independently for each gene with *p*(*m*_*g*_ =1) = 1 − *p*_*m*_ with masking rate = 0.3. The masked input was rescaled by 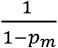 during training to match the expected scale of the unmasked input at inference.

First, the encoder applies a pathway constrained linear projection to obtain the pathway activity representation:

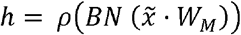

where *W*_*M*_ = *softplus* (*W*_*E*_) ⊙ *M, W*_*E*_ ∈ ℝ^*G*× *P*^ is a learnable encoder weight matrix and *M* ∈ {0,1}^*G*× *P*^ is the Hallmark gene set mask, where *M*_*g*,*j*_ = 1 if gene *g* belongs to pathway *j*. The *softplus* transformation constrains the effective gene to pathway weights to be non-negative, and ⊙ denotes element-wise multiplication. *BN* denotes cell type conditional batch normalization, and *ρ* (·) denotes the GELU^39^ activation function.

The pathway representation is then projected to the latent space:

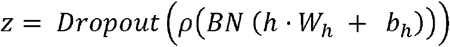

where *W*_*h*_ ∈ ℝ^*G*×*L*^ and *b*_*h*_ ∈ ℝ^*L*^ . The decoder is a multi-layer neural network that maps *z* to the reconstructed gene expression:

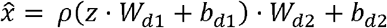

where hidden dimension *H* = 200, *W*_*d*1_ ∈ ℝ^*L*×*H*^, *W*_*d*2_ ∈ ℝ^*H*×*G*^. The pathway dimension was P = 50, corresponding to the Hallmark gene sets, and the latent dimension was *L* =32.

The autoencoder was trained to reconstruct the original gene expression input from the masked 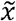 input while jointly optimizing an auxiliary cell type classification objective and a pathway decorrelation penalty:

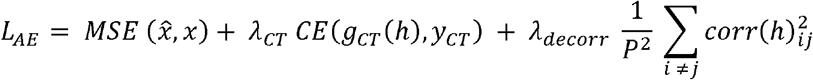

where MSE is the mean-squared-error loss, *g*_*CT*_ (*h*) = *h* · *W*_*CT*_ + *b*_*CT*_ is a linear cell type classification head, is the cross-entropy loss, and *yCT* denotes the known cell type label of each input cell. The matrix *corr* (*h*) ∈ ℝ^*P*×*P*^ denotes the pairwise correlation matrix of pathway activities computed during training. Loss weight hyperparameters are *λ*_*CT*_ = 0.3 and *λ*_*decorr*_ = 0.1 to provide auxiliary regularization for cell type information and pathway decorrelation while maintaining reconstruction loss as the dominant objective during autoencoder training. After training, the autoencoder weights were frozen and used for downstream cell type specific and patient-level prediction.

### Cell type specific classifiers

For each cell type *k*, irAEGIS trains a cell type specific classifier *f*_*k*_ to estimate the patient-level irAE risk score from the pathway representations of cells belonging to that cell type. For patient *p* and cell type *k*, cells were ranked by the L2 magnitude of their pathway activity vector‖ *h* ‖_2_. The aggregated pathway representation *r*_*p*_,_*k*_ ∈ ℝ^*P*^ was computed by averaging the pathway representations of cells in the upper quartile of this ranking.

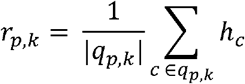

where *q*_*p*_,_*k*_ denotes the upper quartile of cells for patient *p* and cell type *k*. This aggregation strategy emphasizes cells with stronger overall pathway activity, which may contain phenotype signal, rather than diluting this signal by averaging across all cells of the same cell type. The aggregated representation *r*_*p*_,_*k*_ was standardized within each training fold to obtain *u*_*p*_,_*k*_, allowing the L2 regularization penalty to be applied uniformly across pathways with different activation magnitudes.

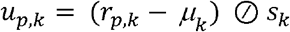

where *µ* ∈ ℝ^*P*^ and *s*_*k*_ ∈ ℝ^*P*^ are the pathway-wise mean and standard deviation of the aggregated representations for cell type *k* among the training patients, and ⊘ denotes element-wise division.

Each cell type specific classifier is an L2 regularized logistic regression model that maps the *p*-dimensional representation *u*_*p*_,_*k*_ to a scalar irAE probability:

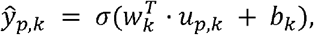

where *w*_*k*_ ∈ ℝ^*P*^, *b*_*k*_ ∈ ℝ^*P*^, *σ*(·) denotes the sigmoid function, and *u*_*p*_,_*k*_ is the standardized aggregated pathway representation *h* for patient *p* and cell type *k*. Each cell type specific classifier was trained by minimizing a class-balanced binary cross entropy loss with L2 regularization.

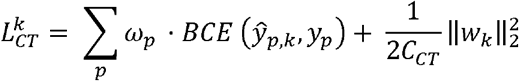

where *y*_*p*_ ∈ {0,1} is the patient irAE label, *ŷ*_*p*_,_*k*_ is the predicted irAE probability for patient *p* using cell type *k*, BCE is the binary cross-entropy loss, *ω*_*p*_ is the class-balanced sample weight for patient p, and *C*_*CT*_ = 0.1 is the inverse regularization parameter. This encourages each cell type specific classifier to identify pathway programs that are predictive within a specific immune cell population while controlling model complexity.

### Patient-level classifier

The patient-level classifier integrates risk scores from informative cell type specific classifiers into a final irAE prediction using a stacked generalization framework^40^. Informative cell types were selected using leave one out cross-validation (LOOCV) on the training patients. Cell types with a training AUC greater than 0.5 were retained as selected informative cell types, denoted by *S*.

To train the patient-level classifier without using predictions from models fit on the same patients, held-out risk scores were generated for each training patient. Specifically, for each selected cell type, the corresponding classifier was trained using all training patients except the held-out patient and then applied to that held-out patient to obtain a predicted irAE probability. These probabilities were transformed to log-odds, using logit transform 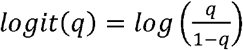, so that the inputs matched the linear log-odds scale of logistic regression. The transformed scores from selected cell types were concatenated to form a patient-level feature vector 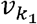.

The patient-level classifier is an L2 regularized logistic regression model that maps *ϕ*_*p*_ to the final irAE probability:

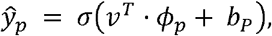

where *b*_*p*_ ∈ ℝ^*P*^, *σ*(·) denotes the sigmoid function, and 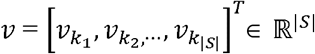 is the learned coefficient vector of the patient-level classifier, and each coefficient *v*_*k*_ corresponds to selected cell type *k* ∈ *S*. Thus *v*_*k*_ quantifies how strongly the out of fold risk score contributes to the final patient level prediction which is derived from the standardized representation *u*_*p*_,_*k*_. The patient-level classifier was trained by minimizing a class-balanced binary cross entropy loss with L2 regularization as follows:

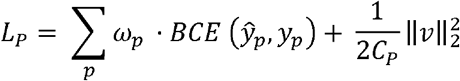

where ∈ ℝ^|*s*|^ is the weight learned by the patient-level classifier, *y*_*p*_ ∈ {0,1} is patient irAE label, *ω*_*p*_ is the class-balanced sample weight for patient p, and *C*_*p*_ = 1 is the inverse regularization parameter. This design allows the classifier to combine complementary signals across informative cell types while limiting overfitting in small patient cohorts.

### Model interpretability

irAEGIS provides model-based explanations at two levels. Dataset-level explanations summarize the cell types, pathways, and genes most associated with differences between irAE positive and irAE negative patients across the dataset. Patient-level explanations summarize how the trained prediction model uses cell type and pathway scores to make a prediction for an individual patient.

### Dataset-level explanations

Cell type importance was quantified using the discriminative value of each cell type specific classifier. Specifically, cell types were ranked by the cross-validation AUC of their corresponding cell type classifiers, with higher AUC indicating greater ability to distinguish the two patient outcome groups.

For pathway-level explanations, differential pathway activity was computed using the raw aggregated pathway representation *r*_*p*,*k*_ which was obtained by aggregating the pathway activity representation *h* of the upper quartile cells from patient *p* and cell type *k*. For pathway *j* in cell type *k*, differential pathway activity was computed as the signed difference between the mean aggregated pathway activity in irAE positive and irAE negative patients.

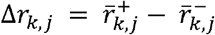

where 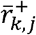 and 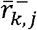 denote the mean aggregated activity of pathway in *j* cell type *k* among irAE positive and irAE negative patients. A positive value indicates higher pathway activity in irAE positive patients, whereas a negative value indicates higher pathway activity in irAE negative patients. Pathways within each cell type were ranked by 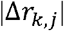.

Gene level attribution scores were obtained by projecting classifier-aligned differential pathway activity back to genes through the Hallmark masked encoder weight matrix *W*_*M*_.

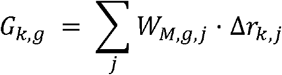

Here, *G*_*k*,*g*_ represents the dataset-level attribution score for gene *g* in cell type *k*. The magnitude |*G*_*k*,*g*_| ranks genes by their cumulative contribution through Hallmark pathway membership, and the sign indicates the direction of association. A positive value indicates stronger association with irAE positive patients, whereas a negative value indicates stronger association with irAE negative patients.

### Patient-level explanations

For an individual patient, irAEGIS uses the fixed weights of the final trained cell type classifiers and patient-level classifier to quantify how selected cell types and pathways contribute to the prediction. For patient *p* and cell type *k*, the cell type specific prediction score before the sigmoid transformation is:

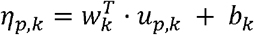

where *u*_*p*_,_*k*_ is the standardized pathway representation for patient *p* and cell type *k, w*_*k*_ is the trained classifier weight vector and *b*_*p*_ is the bias associated with the cell type specific classifier *k*.

The contribution of pathway *j* within cell type *k* is:

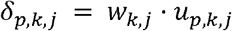

Thus, the cell type specific prediction score can be written as:

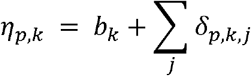

The patient-level classifier then linearly combines the selected informative cell types into the final prediction score before sigmoid transformation:

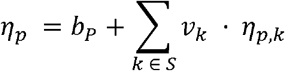

Where *v*_*k*_ is the trained patient-level classifier weight for cell type *k, b*_*p*_ is its bias, and *S* is the set of selected informative cell types.

The pathway importance after patient-level weighting is:

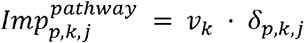

and the patient-level cell type importance is:

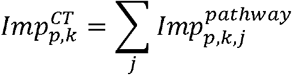

Gene importance scores were computed within each patient and cell type by projecting pathway attributions through the Hallmark masked encoder weights:

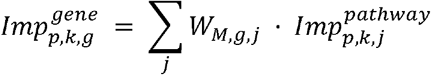

Bias terms are included to compute the full prediction score but are not reported as attributions since these terms are constant for the model. Cell type and pathway contributions are computed from the trained linear classifiers and gene scores are pathway guided attribution scores. These model derived attributions are intended to guide biological interpretation.

### Experimental setup

The training was divided into two stages. In the first stage, the pathway constrained denoising autoencoder was trained using an unsupervised objective, without using patient irAE outcome labels to learn biologically anchored pathway representation from scRNA-seq profiles. The autoencoder is trained to map cells into a low-dimensional pathway activity representation and reconstruct the original gene expression profile. The encoder consists of a pathway-masked linear projection followed by cell type conditional batch normalization and GELU activation. The latent projection included a linear layer, batch normalization, GELU activation, and dropout (p = 0.1). The decoder is a two-layer neural network with hidden dimension 200 and GELU activation. The auxiliary cell type head is a single linear layer mapping the pathway activity representation to cell type class logits. This stage used reconstruction loss, auxiliary cell type classification loss, and pathway decorrelation loss. The autoencoder was trained for 80 epochs using Adam^41^ optimizer with a batch size of 2048 cells, learning rate of 1×10□^3^, and weight decay of 1×10 □^4^.

In the second stage, the encoder and decoder weights were frozen. Cell type specific classifiers were trained independently for each cell type. Then, the patient-level classifier was trained within each cross-validation fold using only the training patients. The cell type specific classifiers and patient-level classifier were trained for a maximum of 2000 iterations with a convergence tolerance of 1×10□^4^. Patient-level irAE prediction was evaluated using LOOCV. In each fold, one patient was held-out for testing, and the remaining *N* = 1 patients were used for training. Within each cross-validation fold, cell type specific classifiers were trained independently for each cell type. Cell types with cross-validation AUC greater than 0.5 on the training patients were retained as informative cell types. The patient-level classifier was then trained using stacked held-out predictions from the retained cell type specific classifiers. The same LOOCV protocol was applied to all real and simulated datasets, and identical fold partitions and preprocessing were used across all methods. All supervised components, including cell type specific classifiers, cell type selection, and the patient-level classifier, were trained using training patients only within each cross-validation fold. In each LOOCV fold, the held-out patient was used for testing. Patient-level performance was assessed using the area under the receiver-operating-characteristic curve (AUC) and the area under the precision-recall curve (AUPRC) computed from the held-out patient predictions aggregated across all LOOCV folds.

To assess the contribution of individual irAEGIS components, we performed component ablations on the four real datasets. Each ablation removed or replaced one design choice while keeping preprocessing, fold partitions, and training settings fixed. We evaluated three variants: removal of the auxiliary cell type classification loss, removal of the pathway decorrelation loss, and the use of fixed Hallmark pathway scores without the autoencoder representation learning. In the fixed Hallmark score ablation, each cell was represented by 50 pathway scores computed as the mean log normalized expression of genes in each Hallmark gene set. Ablation performance was assessed using patient-level AUC with detailed results reported in **Supplementary Table S3**.

We compared irAEGIS with seven baseline methods. These included three DL methods for single-cell phenotype prediction, ScRAT, singleDeep, and Hierarchical MIL, and four traditional machine learning baselines, Cell LR in which a logistic regression classifier was trained at the cell level and then aggregated to the patient-level, as well as Pseudobulk LR, Random Forest, and XGBoost trained on patient-level pseudobulk gene expression profiles. All baselines were evaluated using the same LOOCV fold partitions. For ScRAT, singleDeep, and Hierarchical MIL, default settings from their source repositories were used with repository sources provided in **Supplementary Table S4**. For performance evaluation, irAEGIS was assessed using cross validation. For interpretability analysis, irAEGIS was trained on the full dataset, and explanations were computed using the final trained model with encoder, cell type classifiers, and patient-level classifier weights fixed.

### Evaluation metrics

Given a dataset with *N* patients, ground truth irAE labels *y*_*p*_ ∈{0,1} and predicted probabilities *ŷ*_*p*_ ∈ [0,1] for patients *p* = 1, …, *N*, we used the following metrics to evaluate the prediction performance and interpretability of irAEGIS.

For prediction performance, we reported the area under the receiver operating characteristic curve (AUC), and the area under the precision recall curve (AUPRC) computed on the held-out patients across cross-validation folds. AUC measures the model’s ability to rank irAE positive patients above irAE negative patients across decision thresholds, while AUPRC is informative under class imbalance. Higher AUC and AUPRC indicate better predictive performance.

To evaluate the explanations of irAEGIS, we used three complementary metrics: fidelity score, class contrastivity score, and ground truth recovery.

Fidelity score: The fidelity score is computed as the decrease in patient-level AUC after masking irAEGIS important features. We assessed fidelity at two biological resolutions: pathway-level masking where the genes belonging to irAEGIS’s top 20 important pathways for each cell type are masked, and cell type masking, where the informative cell types with cross-validation classifier AUC > 0.5 are masked. A higher fidelity score indicates that important features attributed by irAEGIS have greater influence on model predictions.

Class contrastivity score: The class contrastivity score measures whether irAEGIS identified distinct pathway programs for irAE positive and irAE negative patients. Within each cell type, important pathways were defined as those whose absolute differential pathway activity exceeded the within cell type mean by more than one standard deviation. For each cell type *k*, the selected pathways were partitioned into irAE positive and irAE negative class. Pathways with positive differential pathway activity were associated with irAE positive class and pathways with negative differential pathway activity were associated with irAE negative class. Let *P*^+^ denote the set of irAE positive pathways, and *p*^*−*^ denote the set of irAE negative pathways. Class contrastivity is computed as 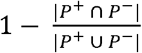. A higher contrastivity score indicates that irAEGIS assigns more distinct pathway programs to the two irAE outcome groups within each cell type.

Ground truth recovery: For simulated datasets, the injected cell type, pathway, and gene signals are known, allowing direct evaluation of whether irAEGIS recovered the ground truth signal. Pathway recovery was evaluated using Recall, defined as the fraction of injected ground truth pathways recovered among the top 5 irAEGIS attributed important pathways for each cell type where signal is injected. Gene recovery was evaluated using Precision, as the fraction of top 30 irAEGIS attributed important genes within each signal carrying cell type that belonged to the injected ground truth gene set. Higher recall and precision indicate better recovery of the injected biological signal.

## Supplementary Tables

**Supplementary Table S1: Predictive performance of irAEGIS and baseline methods**. AUC and AUPRC values are reported for irAEGIS and seven competing methods across all real and simulated datasets.

**Supplementary Table S2: Summary of datasets used in this study**. The table summarizes the real patient cohorts and provides the simulation parameters for the restricted signal and distributed signal settings.

**Supplementary Table S3: Ablation analysis of irAEGIS components**. The table reports AUC values across four real patient datasets for the full irAEGIS model and ablated variants for evaluating the contribution of cell type auxiliary loss, pathway decorrelation loss, and learned autoencoder representation.

**Supplementary Table S4: Baseline method details**. The table provides the source repositories and versions for the baseline deep learning methods.

**Supplementary Table S5: Sample level metadata for datasets used in the study**. The table provides sample level information, including sample identifier, patient identifier, species, tissue type, anatomical site, treatment timepoint, ICI treatment states, irAE status, irAE grade, irAE severity group, and disease group.

## DATA AVAILABILITY

All datasets used for evaluation are publicly available from the NCBI Gene Expression Omnibus (GEO) at https://www.ncbi.nlm.nih.gov/geo/. All data information is illustrated in **Supplementary Table S5**.

## CODE AVAILABILITY

The irAEGIS method is provided as an open-source Python package on GitHub: https://github.com/QSong-github/irAEGIS.

